# Deletion of the voltage-gated calcium channel gene, Ca_V_1.3, reduces Purkinje cell dendritic complexity without altering cerebellar-mediated eyeblink conditioning

**DOI:** 10.1101/2025.03.27.645586

**Authors:** Annette J. Klomp, Martha Pace, Jacqueline B. Mehr, Maria Fernanda Hermosillo Arrieta, Cessily Hayes, Anthony Fleck, Shane Heiney, Aislinn J. Williams

## Abstract

Genetic variation in *CACNA1D*, the gene that encodes the pore-forming subunit of the L-type calcium channel Ca_V_1.3, has been associated with increased risk for neuropsychiatric disorders that display abnormalities in cerebellar structures. We sought to clarify if deletion of Ca_V_1.3 in mice would induce abnormalities in cerebellar cortex cytoarchitecture or synapse morphology. Since Ca_V_1.3 is highly expressed in cerebellar molecular layer interneurons (MLIs) and L-type channels appear to regulate GABA release from MLIs, we hypothesized that loss of Ca_V_1.3 would alter GABAergic synapses between MLIs and Purkinje cells (PCs) without altering MLI numbers or PC structure. As expected, we did not observe changes in the numbers of MLIs or PCs. Surprisingly, Ca_V_1.3 KO mice do have decreased complexity of PC dendritic arbors without differences in the number or structure of GABAergic synapses onto PCs. Loss of Ca_V_1.3 was not associated with impaired acquisition of delay eyeblink conditioning. Therefore, our data suggest that Ca_V_1.3 expression is important for PC structure but does not affect other measures of cerebellar cortex morphology or cerebellar function as assessed by delay eyeblink conditioning.

## Introduction

Genetic variation in *CACNA1D*, the gene that encodes the pore-forming subunit of the L-type calcium channel Ca_V_1.3, has been associated with increased risk for autism spectrum disorder (Fu et al., 2022; Pinggera et al., 2018, 2017, 2015; Pinggera and Striessnig, 2016), bipolar I disorder (Ament et al., 2015; Ross et al., 2016), epilepsy (Pinggera et al., 2017; Pinggera and Striessnig, 2016), schizophrenia (Cross-Disorder Group of the Psychiatric Genomics Consortium, 2013; Pardiñas et al., 2018), hyperaldosteronism (Flanagan et al., 2017; Ortner et al., 2020; Pinggera et al., 2018, 2017, 2015; Scholl et al., 2013; Tan et al., 2017), and deafness and bradycardia (Baig et al., 2011) suggesting that Ca_V_1.3 plays an important role in a wide variety of systems. Schizophrenia, autism spectrum disorder, and bipolar disorder subjects display abnormalities in cerebellar volume, gene expression, connectivity patterns of cerebellar circuits, and cerebellar-dependent motor and cognitive behaviors (Andreasen et al., 1997; Andreasen and Pierson, 2008; Becker and Stoodley, 2013, 2013; Bolbecker, 2009; Crespo-Facorro et al., 2007; Guidotti, 2000; Johnson, 2018; Levitt et al., 1999; Lundin et al., 2021; Parker et al., 2014; Shinn et al., 2017; Stoodley et al., 2017). However, the cellular mechanisms that drive changes in the cerebella of patients with these neuropsychiatric disorders are still unknown. Given the genetic links between L-type calcium channels and neuropsychiatric disorders, a better understanding of the role of L-type channels in cerebellar microcircuit structure may be broadly applicable to our understanding of neuropsychiatric disorders.

One possible role for L-type calcium channels in the cerebellum is in modulating the development or function of the cerebellar cortex microcircuit. In the cerebellar cortex microcircuit, PCs receive excitatory input from climbing fibers from the inferior olive and the parallel fibers of granule cells, as well as inhibitory signals from MLIs (Eccles et al., 1966a, 1966b). PCs then send inhibitory signals to the deep cerebellar nuclei to trigger initiation of movement, thought, or behaviors (Green and Steinmetz, 2005). MLIs consist of two subsets of neurons: basket cells (BCs) which generally reside in the basal one-third of the molecular layer, and stellate cells (SCs) which typically reside in the apical two-thirds of the molecular layer. BC axons wrap around and synapse on the initial segment of the PC axon via a structure termed a pinceau (French for “brush”), (Cajal, 1911), whereas SCs are generally located more superficially in the molecular layer and synapse onto the dendrites of PCs (Eccles et al., 1966b).

Several cerebellar cell types, such as PCs, Golgi cells, and MLIs, express *Cacna1d* mRNA according to mouse single-nucleus transcriptomic datasets (Kozareva et al., 2021; Saunders et al., 2018) and immunohistochemical and proteomic data (Hell, 1993; Uhlén et al., 2015). L-type channel activity has been detected in developing immature PCs but is less apparent in mature PCs (Gruol et al., 2006; Tringham et al., 2007), suggesting that Ca_V_1.3 in PCs may be important primarily in development. L-type currents have been detected in MLIs and are thought to modulate GABA release from MLIs to PCs (Rey et al., 2020), suggesting a potential role for Ca_V_1.3 in mature MLI function.

We explored whether ubiquitous germline deletion of Ca_V_1.3 results in abnormal morphological features in PCs and MLIs, and whether this alters cerebellum-dependent learning using the delay eyeblink conditioning paradigm. Given the high expression and electrophysiological role of Ca_V_1.3 in cerebellar MLIs, we predicted that loss of Ca_V_1.3 would alter MLIs and the GABAergic synapse between MLIs and PCs. However, we have instead found that Ca_V_1.3 deletion alters PC morphology without appearing to affect the synaptic structure or fate specification of MLIs, and without impacting acquisition of cerebellum-dependent delay eyeblink conditioning.

## Methods

### Mice

The generation of Ca_V_1.3 knockout (KO) mice (Cacna1d^tm1Jst^) has been described previously (Lauffer et al., 2022; Platzer et al., 2000). Breeding pairs of Ca_V_1.3^+/-^ mice were maintained on a C57BL/6NTac background by crossing Ca_V_1.3^+/-^ offspring with C57BL/6NTac wild-type (WT) mice purchased from Taconic Biosciences (Rensselaer, NY). To generate experimental animals, Ca_V_1.3^+/-^ males were bred to Ca_V_1.3^+/-^ females to obtain male and female Ca_V_1.3 WT and KO littermates. All mice were adults (at least 10 weeks old) at the time of use. Sample sizes are indicated in each figure. All experiments were carried out in a manner to minimize pain and discomfort. All experiments were conducted according to the National Institute of Health guidelines for animal care and were approved by the Institutional Animal Care and Use Committee at University of Iowa.

### Histology

Ca_V_1.3 KO and wild type littermates 13-30 weeks old (*n*=6-8 per genotype) were anesthetized with 17.5mg/ml Ketamine / 2.5mg/ml Xylazine at a dose of 0.1ml per 20g and perfused with 4% paraformaldehyde in 0.1M phosphate buffer (PB). Whole brains were dissected and immersed in 30% sucrose for 72 hours. Brains were rinsed in PB and frozen in optimal cutting temperature compound. Brain tissue was sectioned on a cryostat into 20-μm-thick sagittal sections, mounted on slides, and stored at −20°C until use. All histology was performed in cerebellar vermis.

### Immunostaining

Blocking buffer was made from normal donkey serum (5%) and Triton-X (0.1%) in PB. Primary antibodies used included: rabbit anti-Calbindin D28K (ThermoFisher, Cat: 711-443) 1:250, mouse anti-parvalbumin (Swant, Cat: PV 235) 1:250, rabbit anti-HCN1 (Synaptic Systems, Cat: 338-003) 1:50, mouse anti-GAD6 (DSHB, Cat: AB_528264) 1:250, and mouse anti-Aldolase C 4A9 (Novus Bio, Cat: NBP2-25145) 1:500. All primary antibodies were incubated on sections overnight at 4°C. Secondary antibodies used included donkey anti-rabbit 488 (Jackson, Cat: 711-545-152) 1:500 and donkey anti-mouse 594 (Jackson, Cat: 715-585-151) 1:500. Secondaries for Aldolase C labeling were incubated overnight at 4°C and all other secondaries were incubated at RT for 2-4 hours. Sections were incubated with DAPI Solution (Thermo Scientific) 1:1000 for 1 minute and coverslips were mounted with Prolong Diamond Antifade Mountant (Invitrogen).

### Golgi Staining

Ca_V_1.3 KO and littermate wild type mice at least 10 weeks old (*n*=6-8 per genotype) were anesthetized with isoflurane. Whole brains were dissected and stained with the FD Rapid GolgiStain Kit (FD NeuroTechnologies). Brains were immersed in impregnation solution A/B for 2 weeks then immersed in solution C for 5 days before sectioning at 100μm and staining sections with staining solution D/E per manufacturer protocol. Sections were dehydrated with ethanol and coverslips were mounted with Prolong Diamond Antifade Mountant (Invitrogen).

### Microscopy

Sections were imaged either at 20x on an Olympus IX83 fluorescence microscope and stitched together using Olympus CellSens Dimension 2.3 software or at 40x in Z-stacks on a Leica SPE Confocal Microscope. Analysis was performed with ImageJ or Imaris 9.9.1. Filament tracing was done with the Imaris Filaments tool. For dendrite tracing, Imaris settings were set to largest diameter of 10μm and thinnest diameter of 1μm. To detect dendritic spines, Imaris settings were set to thinnest diameter spine head of 0.3μm and max spine length of 4μm. Spine classifications were made using the standard default settings in Imaris. For GABAergic synapses, Imaris settings were set to thinnest diameter spine head of 0.3μm and max spine length of 1μm.

### Eyeblink Conditioning

Delay eyeblink conditioning was performed as previously described (Heiney et al., 2018, 2014). Briefly, mice were implanted with a titanium headplate and habituated to head fixation on a treadmill for 2 days. During habituation, no stimuli were presented. Training was performed for 9 days and consisted of 100 trials/day, with 90 CS-US paired trials and10 conditioned stimulus (CS)-alone trials pseudorandomly interleaved. Mice were free to walk on the treadmill during the entire session. The CS was an LED, and the unconditioned stimulus (US) was a puff of air (20 PSI source pressure) of 20-30 ms duration directed at the cornea via a 23-gauge needle placed 3 mm from the mouse’s eye. These pulse durations resulted in 6-8 PSI puff intensities measured at the end of the needle. Interstimulus interval was set at 250 ms. CS-alone trials were included to assess conditioned responses (CRs) uncontaminated by the US or unconditioned response (UR). Trials were separated by a variable inter-trial interval (ITI) that averaged 15-25 s.

### Statistical Analysis

Data were graphed and analyzed using GraphPad Prism 9.0 (GraphPad Software, San Diego, CA) and R (R 4.1.1, emmeans 1.7.0, lme4 1.1.27.1, lmerTest 3.1-3, effectsize 0.5), except for eyeblink conditioning figures which were generated in Adobe Illustrator from MATLAB plots. Data are graphically represented as mean ± standard error of the mean (SEM) for each group. Data were analyzed using the statistical test noted in results (linear mixed model, two-way repeated measures ANOVA, one-way ANOVA, or Student’s *t*-tests with appropriate follow-up testing). Behavioral data from Ca_V_1.3 KO mice do not show strong sex by genotype interaction effects (Lauffer et al., 2022; McKinney and Murphy, 2006), so we have combined sexes for histological experiments. Eyeblink conditioning data were analyzed and reported in the results including sex as a variable, but since no sex differences were observed, results are graphed with males and females combined. Results were considered significant when p<0.05 (denoted in all graphs as follows: *p<0.05; **p<0.01).

## Results

### Loss of Ca_V_1.3 does not alter cerebellar cortex thickness

Given the high expression of *Cacna1d* in cerebellar MLIs (Kozareva et al., 2021; Saunders et al., 2018) and known roles for L-type channels in neuronal structure in hippocampal neurons (Kim et al., 2017; Stanika et al., 2016, 2015), we hypothesized that Ca_V_1.3 is important for normal cerebellar morphology. We first sought to determine whether Ca_V_1.3 KO mice display abnormal cerebellar cortex layer thickness (Figure 1a). We observed no Ca_V_1.3-dependent differences in granular layer thickness in any lobule (n=8/genotype, 2-way ANOVA, main genotype effect, F_1,109_=2.15, p=0.15, genotype x lobule interaction effect, F_7,109_=1.89, p=0.08) (Figure 1b) or molecular layer thickness in any lobule (n=8/genotype, 2-way ANOVA, main genotype effect, F_1,109_=0.50, p=0.48, genotype x lobule interaction effect, F_7,109_=1.56, p=0.15) (Figure 1c). We did observe granular and molecular layer thickness differences between lobules that were independent of genotype (granular layer thickness; main effect lobule, F_7,109_=30.51, p<0.01; molecular layer thickness; main effect lobule, F_7,109_=5.09, p<0.01) (Figure 1b-c). Overall, deletion of Ca_V_1.3 does not appear to alter cerebellar granular or molecular layer thickness.

**Fig 1.**
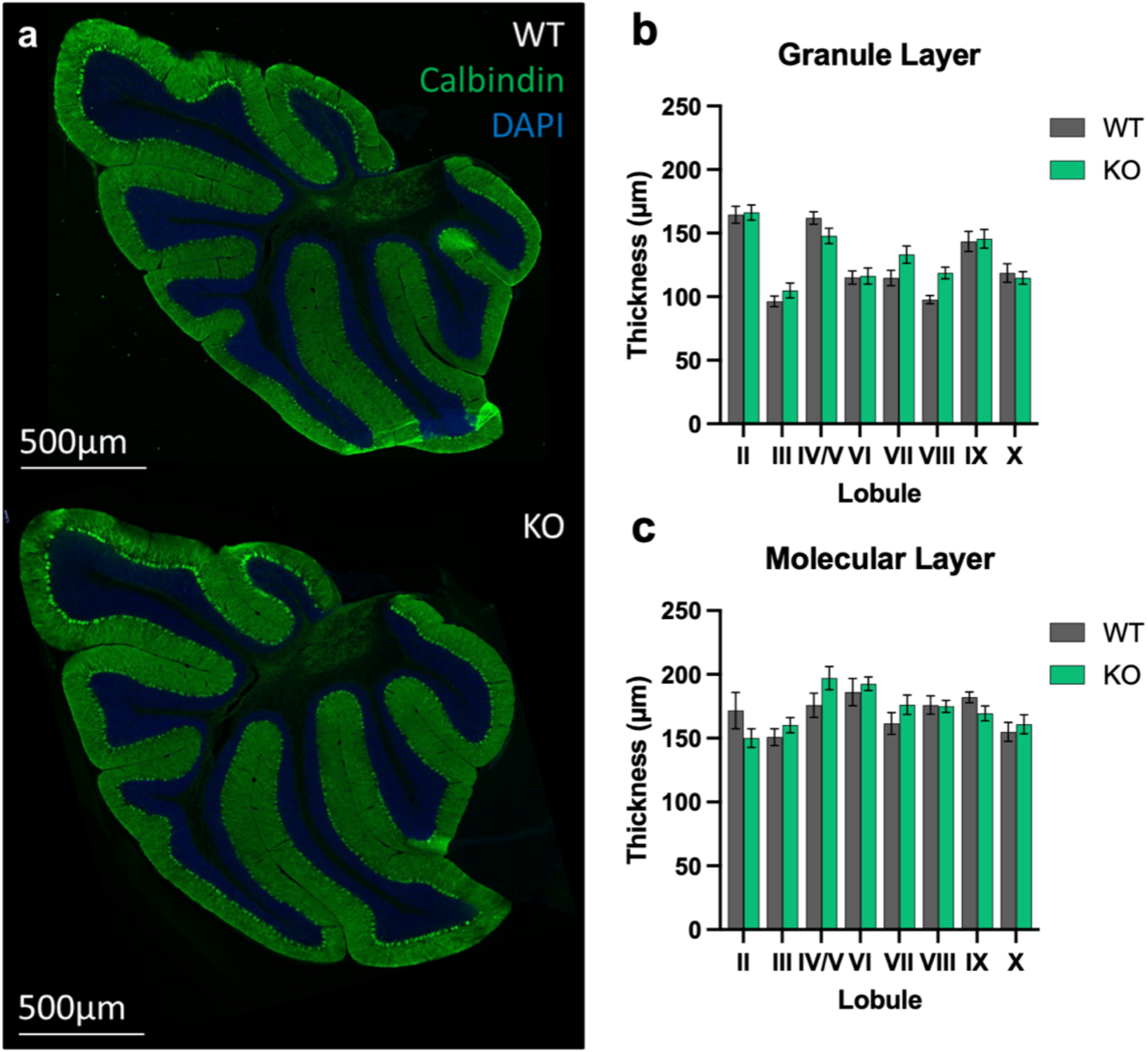
Cerebellar cortex layer thickness varies by lobule. (**a**) Immunofluorescence labeling of calbindin (green) and DAPI (blue) in sagittal sections of cerebellar vermis in Ca_V_1.3 KO and WT littermates. Although cortical layer thicknesses vary across lobules, deletion of Ca_V_1.3 does not alter granule layer (**b**) or molecular layer (**c**) thickness. Data are expressed as mean ± s.e.m.

### Loss of Ca_V_1.3 does not alter PC or MLI density

We next sought to determine whether Ca_V_1.3 KO mice display abnormal numbers of PCs or MLIs. We observed no genotype-dependent differences in numbers of PCs (n=8/genotype, 2-way ANOVA, main genotype effect, F_1,120_=1.55, p=0.22, main lobule effect, F_7,120_=0.52, p=0.82, genotype x lobule interaction effect, F_7,120_=0.19, p=0.99) (Figure 2a) or MLIs (n=6/genotype, DAPI, Linear Mixed Effects Model, main genotype effect, F_1,11_=2.13, p=0.17, genotype x lobule interaction effect, F_7,71_=0.65, p=0.71; PV, Linear Mixed Effects Model, main genotype effect, F_1,7_=0.01, p=0.92, genotype x lobule interaction effect, F_7,66_=1.48, p=0.19) (Figure 2b) in any lobule of the vermis. As with granule and molecular layer thickness, we did observe lobule-dependent differences in MLI density that were independent of genotype (DAPI; main lobule effect, F_7,71_=18.34, p<0.01; PV; main lobule effect, F_7,66_=20.74, p<0.01) (Figure 2b).

**Fig 2.**
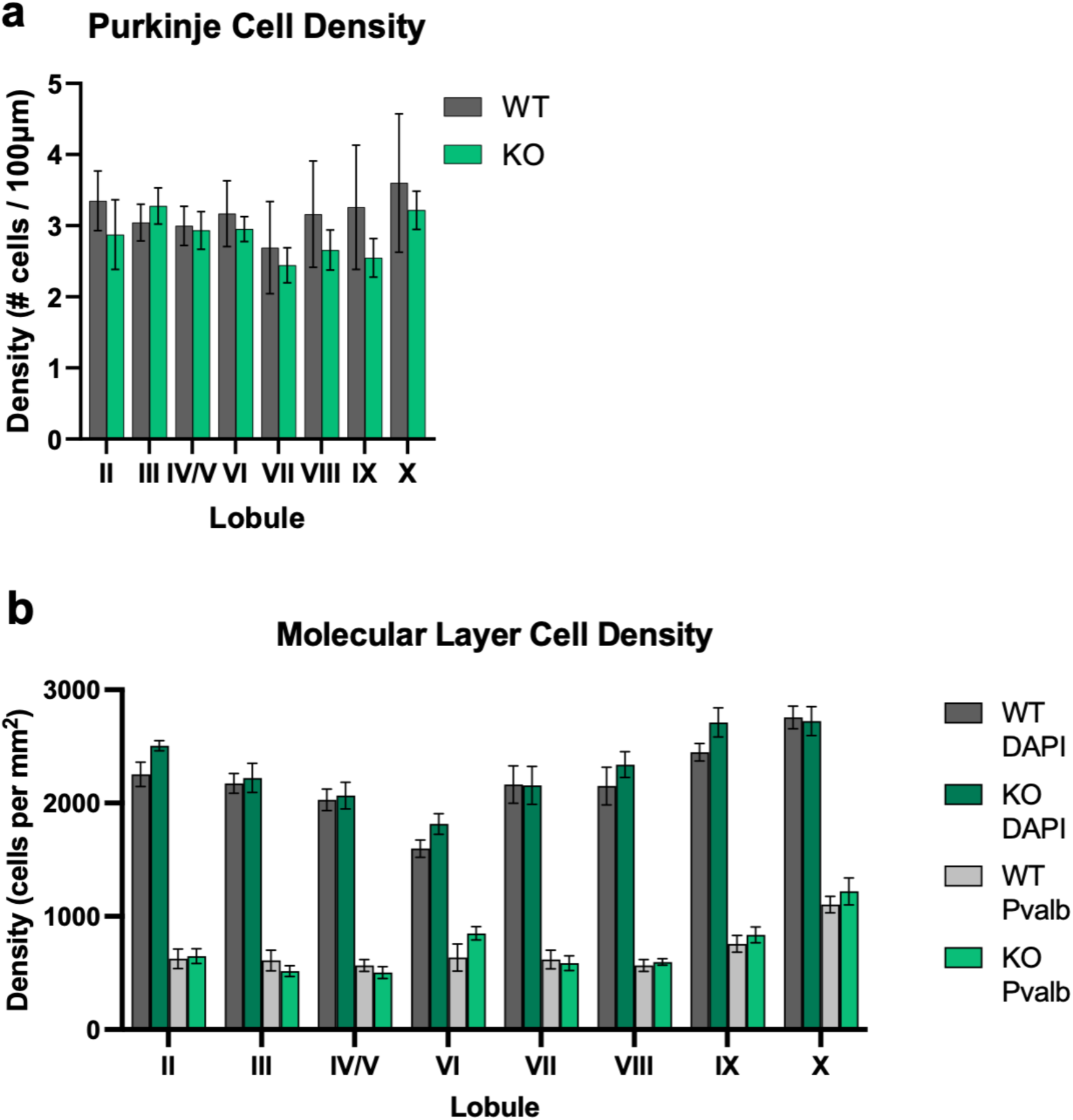
MLI density varies by lobule. Deletion of Ca_V_1.3 does not alter PC density (**a**) or MLI density (**b**), although MLI density does vary significantly as a function of lobule. Data are expressed as mean ± s.e.m.

### Loss of Ca_V_1.3 reduces complexity of PC dendrites

We analyzed PC arborization complexity using a standard Sholl analysis on PCs across the vermis labeled using the Golgi method (Figure 3a). We observed that loss of Ca_V_1.3 resulted in fewer intersections when compared to WT, resulting in a significant reduction in the area under the curve when counting all intersections (n_WT_=13PCs/3mice & n_KO_=16PCs/3mice, nested t test, main genotype effect, F_1,27_=4.22, p<0.05) (Figure 3b). To determine where differences appeared in dendritic arbors, we divided the arbors into 2μm bins (Figure 3c), which showed that Ca_V_1.3-dependent differences in PC dendritic arborization appear to be driven primarily by the distal arbors (n_WT_=13PCs/3mice & n_KO_=16PCs/3mice, Linear Mixed Effects Model, main genotype effect, F_1,27_=4.15, p=0.05, main radius effect, F_115,3105_=50.83, p<0.01, genotype x radius interaction effect, F_115,3105_=2.77, p<0.01, Estimated Marginal Means, p<0.01 at 90μm, 100-140μm, and 150μm, p<0.05 from 100-142μm, 146-160μm, 164μm, and 168μm).

**Fig 3.**
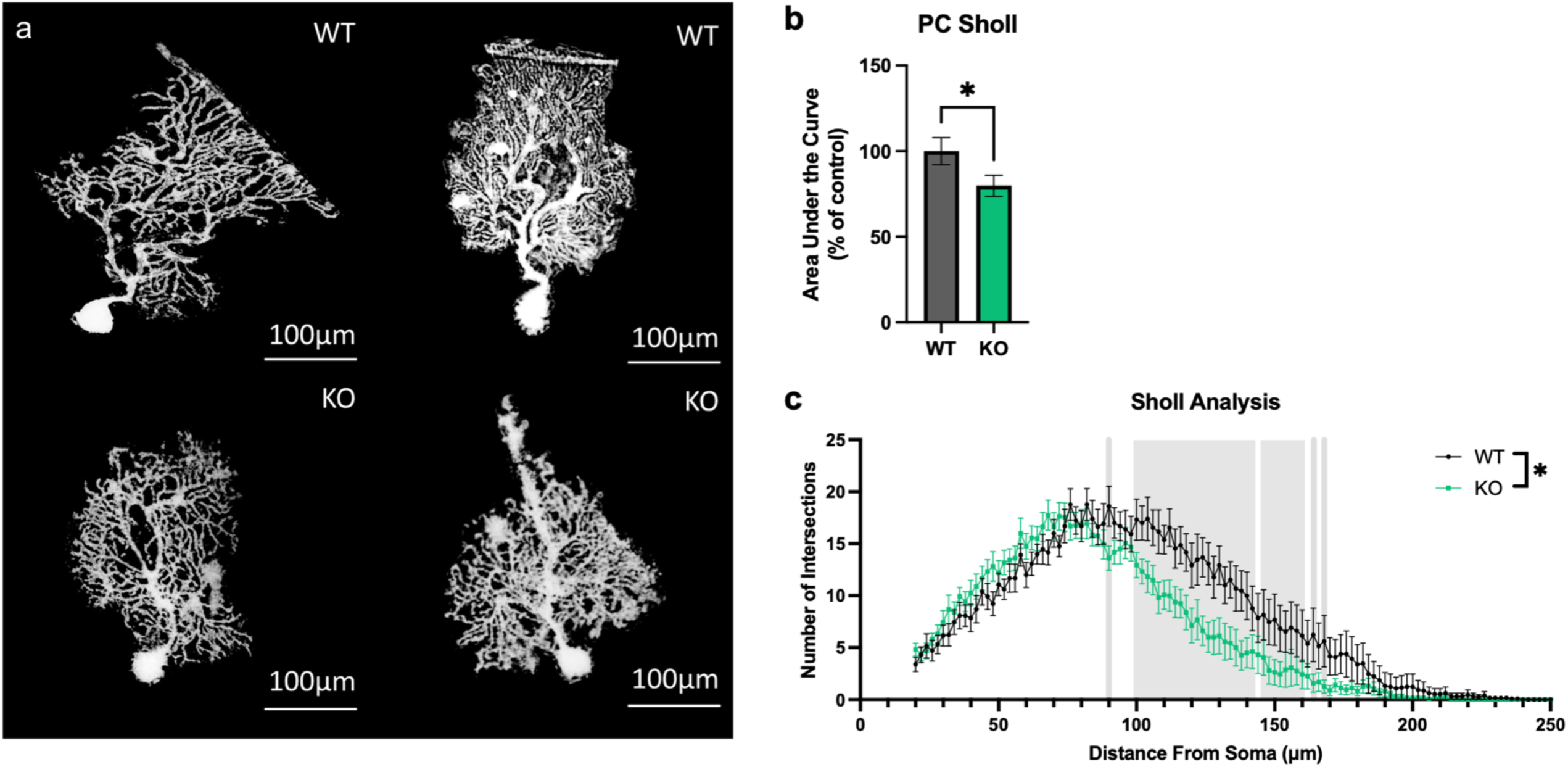
Deletion of Ca_V_1.3 results in less complex distal PC dendritic arborization. (**a**) Golgi staining of PCs from Ca_V_1.3 KO and WT mice from sagittal sections. Deletion of Ca_V_1.3 results in less complex PC dendritic arborization (**b**) driven primarily by differences in the distal dendrites (**c**). Distances with significant differences (*p<0.05) in number of intersections between Ca_V_1.3 KO and WT are highlighted in grey. Data are expressed as mean ± s.e.m.

Given that changes in PC dendritic spines and arborization have been associated with impairments in cerebellar-dependent behaviors (Kloth et al., 2015), we next sought to determine whether Ca_V_1.3 KO mice display abnormal PC dendritic spines or dendritic arborization (Figure 4a). We found that spine density varied by lobule (n=5/genotype, Linear Mixed Effects Model, spine density, main lobule effect, F_7,56_=4.37, p<0.01) (Figure 4b), as did spine classification (n=5/genotype, Linear Mixed Effects Model, main lobule effect, F_7,248_=7.09, p<0.01, main spine type effect, F_7,248_=389.77, p<0.01, lobule x spine type interaction effect, F_21,248_=2.84, p<0.01) (Figure 4c). We observed no Ca_V_1.3-dependent differences in PC spine density (n=5/genotype, Linear Mixed Effects Model, main genotype effect, F_1,8_=0.78, p=0.40, genotype x lobule interaction effect, F_7,56_=0.93, p=0.49) (Figure 4b) nor in distribution of spine classification in any lobule of the vermis (n=5/genotype, Linear Mixed Effects Model, main genotype effect, F_1,8_=0.78, p=0.40, genotype x lobule interaction effect, F_7,248_=1.51, p=0.16, genotype x spine type interaction effect, F_3,248_=0.73, p=0.54, genotype x lobule x spine type interaction effect, F_21,248_=0.28, p=1.00) (Figure 4c).

**Fig 4.**
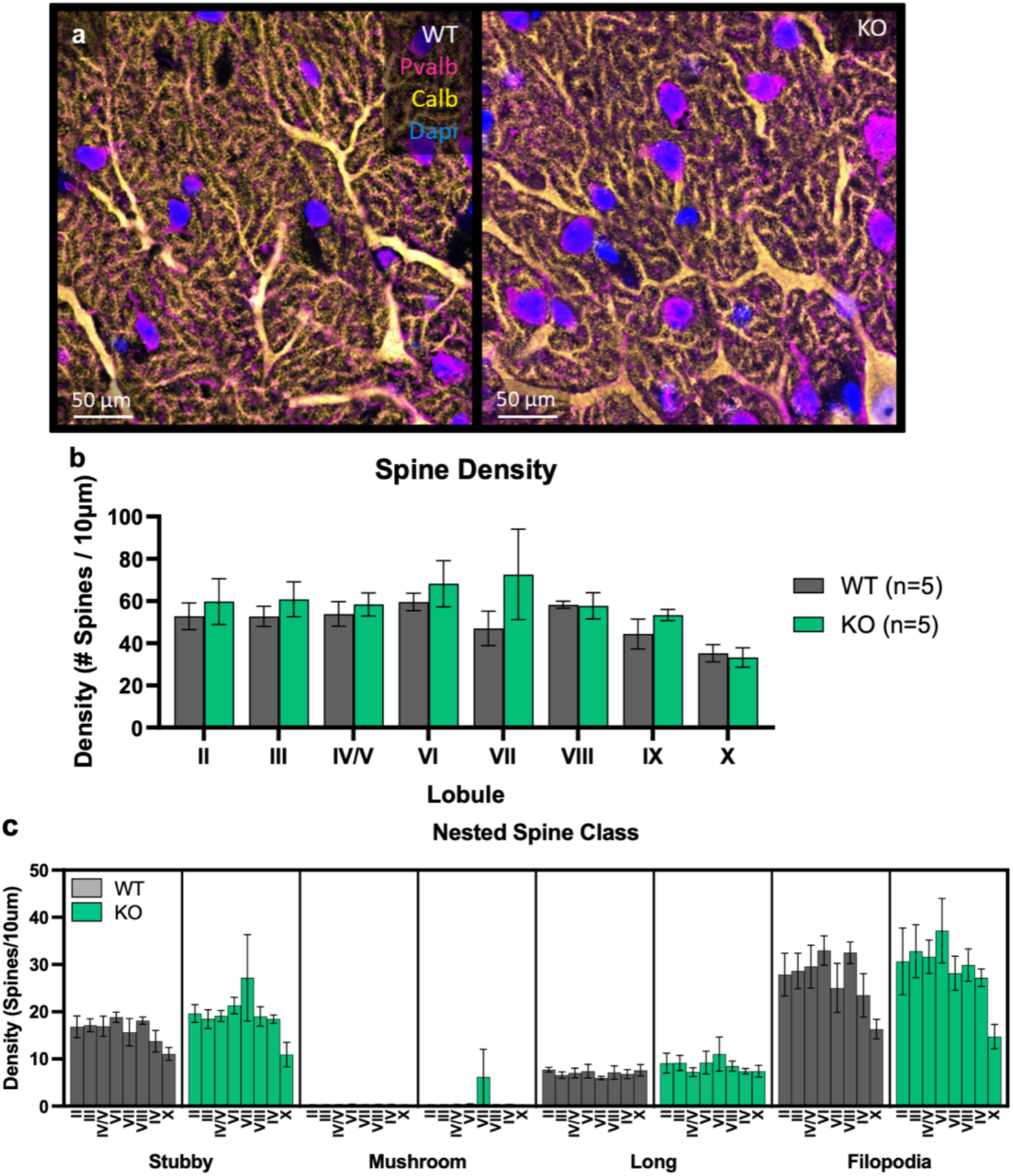
PC spine density varies by lobule. (**a**) Immunofluorescent staining of PV (magenta), calbindin (gold), and DAPI (blue) from Ca_V_1.3 KO and WT mice from sagittal sections. Although spine density varies significantly by lobule, deletion of Ca_V_1.3 does not alter PC spine density (**b**) nor PC spine classification density (**c**). Data are expressed as mean ± s.e.m.

### Loss of Ca_V_1.3 does not alter MLI-PC synapse morphology

Given the high expression of *Cacna1d* in MLIs, we next sought to determine whether Ca_V_1.3 KO MLIs display abnormal presynaptic structures. We first looked at GABAergic synapse density on PC dendrites as measured by GAD6 puncta on calbindin-positive dendrites (Figure 5a). We observed no differences in GABAergic synapse density on PC dendrites (n=4/genotype, Linear Mixed Effects Model, main genotype effect, F_1,6_=0.03, p=0.88, main lobule effect, F_7,39_=0.58, p=0.77, genotype x lobule interaction effect, F_7,39_=1.05, p=0.41) (Figure 5b). We then sought to determine whether Ca_V_1.3 KO mice display abnormal BC pinceau size (Figure 5c). The size of BC pinceaux is typically distributed into zonal modules and this zonal patterning respects PC zonal boundaries (Zhou et al., 2020). Specifically, BC pinceaux are smaller in zebrin II-positive PC zones and larger in PLCβ4-positive and NFH-positive PC zones (Zhou et al., 2020). We replicated this general result (n=5/genotype, Linear Mixed Effects Model, main zebrin effect, F_1,105_=4.05, p<0.05), although this pattern was not observed in lobules IX and X (Figure 5d). We also observed that BC pinceau size varies by lobule (main lobule effect, F_7,105_=2.97, p<0.01). We observed no main effect of genotype in BC pinceau size between Ca_V_1.3 KO and WT mice controlling for PC zones (n=5/genotype, Linear Mixed Effects Model, main genotype effect, F_1,8_=0.17, p=0.69) although there were some trend level genotype interaction effects that did not meet criteria for statistical significance (genotype x lobule interaction effect, F_7,105_=1.78, p=0.10, genotype x zebrin interaction effect, F_1,105_=0.01, p=0.94, lobule x zebrin interaction effect, F_7,105_=2.05, p=0.06, genotype x lobule x zebrin interaction effect, F_7,105_=1.75, p=0.10) (Figure 5d).

**Fig 5.**
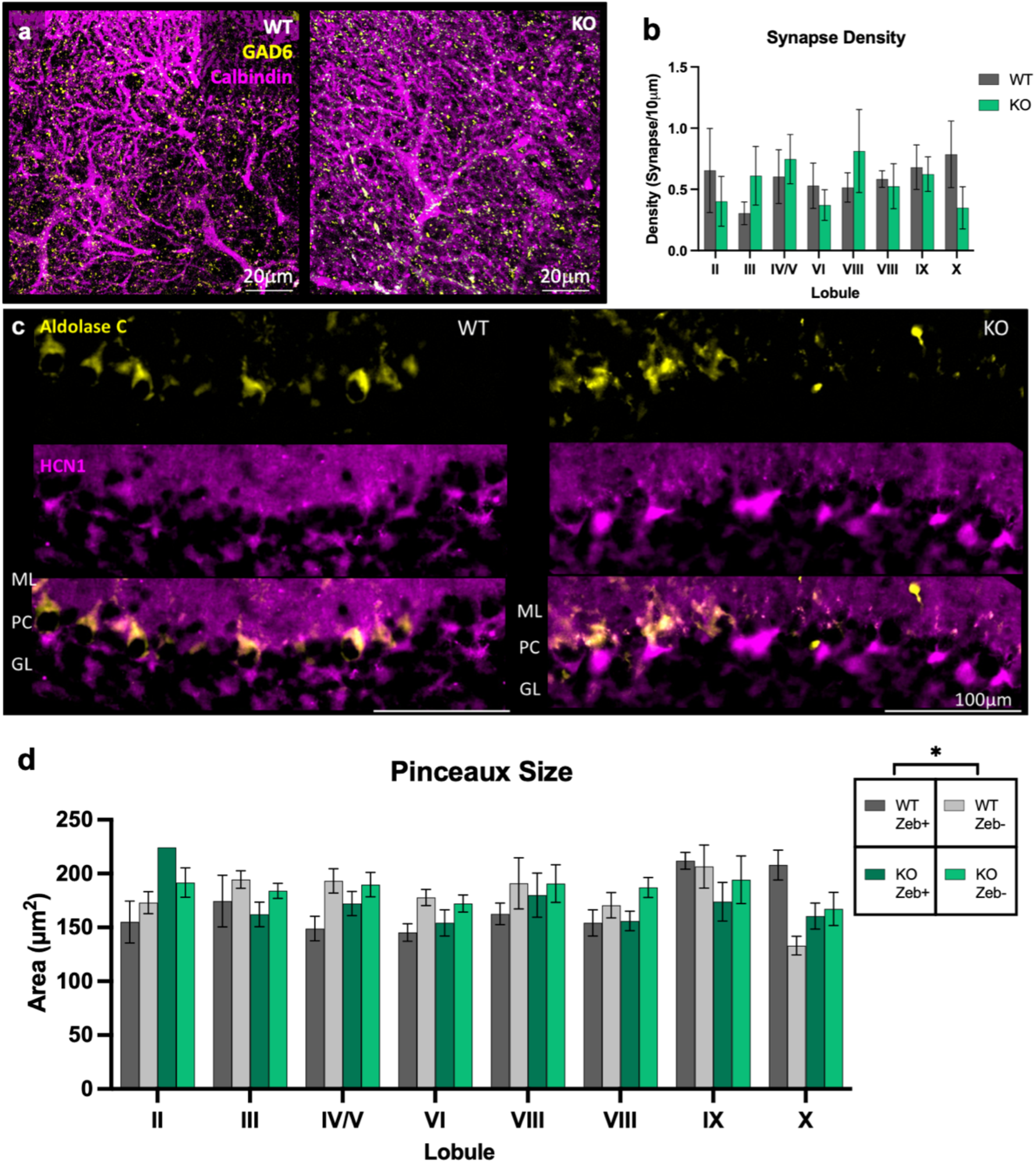
Pinceaux size varies by lobule and zebrin striping. (**a**) Immunofluorescent staining of calbindin (magenta) and GAD6 (gold) from sagittal sections of Ca_V_1.3 KO and WT mice. (**b**) Deletion of Ca_V_1.3 does not alter GABAergic synapse density in the cerebellar molecular layer. (**c**) Immunofluorescent staining of HCN1 (magenta) and aldolase C (gold) from sagittal sections of Ca_V_1.3 KO and WT mice. (**d**) Deletion of Ca_V_1.3 does not alter pinceaux size in the cerebellar molecular layer. Data are expressed as mean ± s.e.m.

### Global deletion of Ca_V_1.3 does not alter delay eyeblink conditioning

Given the reduced dendritic complexity in Ca_V_1.3 KO PCs (Fig. 4a), and the importance of PC function in eyeblink conditioning (Chen et al., 1996; Green and Steinmetz, 2005; Halverson et al., 2015), we examined whether deletion of Ca_V_1.3 altered cerebellum-dependent delay eyeblink conditioning. We examined the learning curves for amplitude of eyelid closure, also called the conditioned response (CR), and observed no effects of genotype or sex and no interaction effect (two-way RM ANOVA, genotype effect F_8,152_=0.53, p=0.83, sex effect F_8,152_=1.6, p=0.13, genotype x sex interaction effect, F_8,152_=1.25, p=0.27). We did find a main effect of session which shows that mice learned the task (F_8,152_=75.98, p<0.01) (Fig. 6a). We then looked at the learning curves for CR percentage, another measure of how learning occurred over the course of training. We again observed no differences between WT and Ca_V_1.3 KO mice (two-way RM ANOVA, genotype effect F_8,152_=0.42, p=0.91, sex effect F_8,152_=0.35, p=0.95, genotype x sex interaction effect, F_8,152_=1.45, p=0.18) although there was a main effect of session (F_8,152_=92.64, p<0.01) (Fig. 6b). Looking at just the final day of training, we saw no group differences for CRs in terms of amplitude (two-way ANOVA, main genotype effect, F_1,37_=0.14, p=0.71, main sex effect, F_1,37_=0.0, p=0.98, genotype x sex interaction effect, F_1,37_=0.07, p=0.80), frequency (two-way ANOVA, main genotype effect, F_1,37_=1.72, p=0.20, main sex effect, F_1,37_=1.71, p=0.20, genotype x sex interaction effect, F_1,37_=0.79, p=0.38), or timing (two-way ANOVA, main genotype effect, F_1,35_=1.43, p=0.24, main sex effect, F_1,35_=1.63, p=0.21, genotype x sex interaction effect, F_1,35_=0.02, p=0.89) (Fig. 6c).

**Fig 6.**
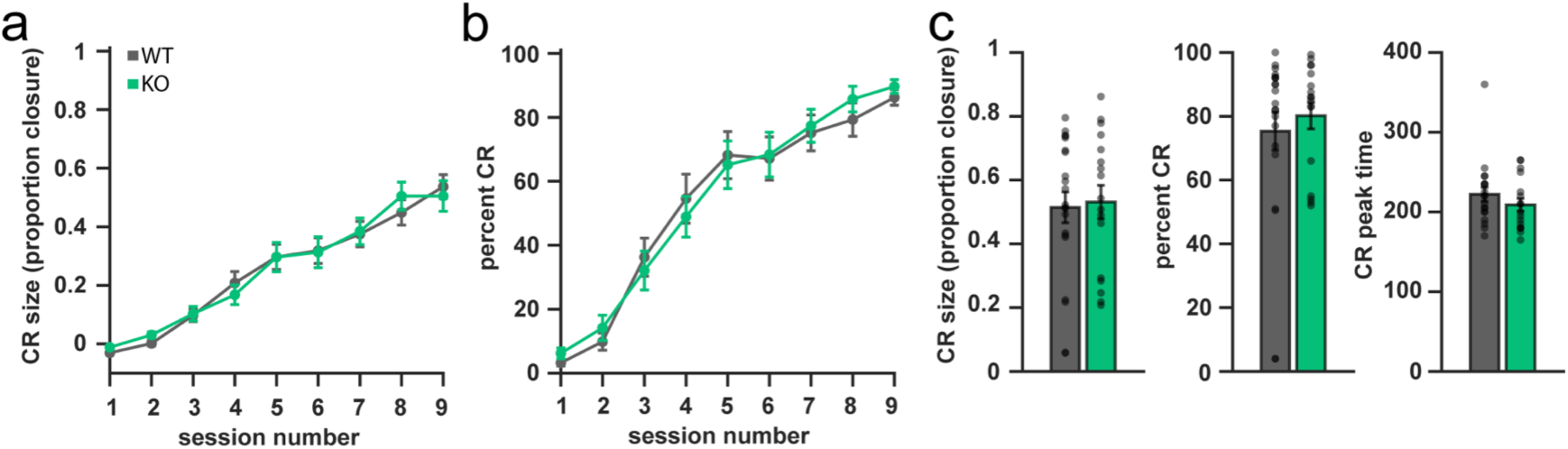
Deletion of Ca_V_1.3 does not impact acquisition of delay eyeblink conditioning. (**a**) No group differences were detected in the learning curves for degree of eye closure during delay eyeblink conditioning training. (**b**) No group differences were detected in the learning curves for conditioned responses during delay eyeblink conditioning training. (**c**) Deletion of Ca_V_1.3 does not alter size of conditioned eyeblink responses (CR size), frequency of conditioned eyeblink responses (percent CR), or timing of conditioned eyeblink responses. Data are expressed as mean ± s.e.m.

## Discussion

Previous work with Ca_V_1.3 KO mice has shown that loss of Ca_V_1.3 results in decreased volume of the auditory brainstem and degeneration of cochlear hair cells without evidence of cerebellar degeneration or atrophy (Hirtz et al., 2011). Consistent with previously published data, our results in general show that loss of Ca_V_1.3 does not alter gross cerebellar structure, particularly with respect to cerebellar cortex thickness and neuronal density. We hypothesized that a loss of Ca_V_1.3 would result in abnormal MLI morphology and alter GABAergic MLI-PC synapses. To our surprise, Ca_V_1.3 deletion alters PC morphology without appearing to affect numbers of PCs or MLIs, or the synaptic structure of MLI inputs onto PCs.

Ca_V_1.3 is crucial for the survival of several neuronal cell types including cochlear hair cells (Hirtz et al., 2011) and hippocampal neurons (Kim et al., 2017; Marschallinger et al., 2015). It is unknown if Ca_V_1.3 is involved in the survival of cerebellar cortex neurons; however, one study of Ca_V_1.3 KO mice suggests that loss of Ca_V_1.3 does not negatively impact gross cerebellar structure (Hirtz et al., 2011). Our results expand upon this previous study through our careful quantification of PCs and MLIs, finding that Ca_V_1.3 is not essential for normal numbers of these neuronal types. We also did not observe impacts of Ca_V_1.3 on the cross-sectional thickness of the granule and molecular layers. We did not attempt to specifically quantify other cerebellar cortex cell types, such as Bergmann glia, or other less abundant types of interneurons such as Golgi cells, Lugaro cells, and unipolar brush cells. Of these other cerebellar cortex cell types, only Golgi cells have high expression of *Cacna1d* mRNA (Kozareva et al., 2021), but we do not know whether there is electrophysiological evidence of L-type activity in Golgi cells. For this reason, we did not prioritize examining them here. In the future, it may be worthwhile to determine whether Golgi cells are impacted by loss of Ca_V_1.3 if evidence of L-type activity in Golgi cells emerges. We also did not examine neurons of the deep cerebellar nuclei, although some of these neurons likely do express Ca_V_1.3. Future studies may also reveal important Ca_V_1.3-dependent differences in these neurons.

L-type channels in the brain are most often thought to act primarily postsynaptically, regulating the expression patterns of membrane receptors and the morphology of dendritic spines in an activity-dependent manner (Shah et al., 2010; Stanika, 2016; Stanika et al., 2015). Ca_V_1.3 interacts with the PDZ domain of proteins that regulate dendritic spine growth and stability. Hippocampal neurons expressing a mutant Ca_V_1.3 channel lacking this PDZ binding domain display an increase in spine elongation (Stanika, 2016). Loss of Ca_V_1.3 has also been shown to increase dendritic spine complexity and reduce cell body size in auditory brainstem neurons (Hirtz et al., 2011). Our data show that there are no differences in dendritic spines on PCs but there is decreased complexity in the distal dendritic arbors of PCs in Ca_V_1.3 KO mice. It is unclear if this change occurs via a cell-autonomous mechanism (related to intrinsic PC developmental programs) or a cell non-autonomous mechanism (how other cells interact with PCs). Future studies could differentiate between these two possibilities by utilizing conditional KO lines targeting either PCs or neurons that synapse on PCs such as granule cells, Golgi cells, and MLIs. We did not measure arborization or dendritic spines of MLIs but given that our results suggest that loss of Ca_V_1.3 alters PC dendritic arborization, Ca_V_1.3 may play a similar role in MLIs or other neuron classes. Future studies may also reveal important Ca_V_1.3-dependent differences in dendrites of other neurons.

In addition to their roles in postsynaptic function, recent work has also found that L-type channels have presynaptic functions in some cell types (Dolphin and Lee, 2020). For example, in cochlear inner hair cells, Ca_V_1.3 is expressed at the presynaptic ribbon, and influx of Ca^2+^ through Ca_V_1.3 allows for fast and sustained glutamate release (Brandt et al., 2003; Platzer et al., 2000). Presynaptic Ca_V_1 channels in hippocampal interneurons and cerebellar MLIs modulate short-term plasticity via regulation of GABA release (Jensen and Mody, 2001; Rey et al., 2020; Stanika et al., 2016). Additionally, loss of Ca_V_1.3 alters presynaptic bouton size in the hippocampus (Kim et al., 2017), suggesting that Ca_V_1.3 has structural as well as electrophysiological presynaptic functions in this structure. However, we did not find any differences in measures of GABAergic MLI-PC synapse number or basket cell pinceau size. These data suggest that Ca_V_1.3 does not regulate presynaptic structure in cerebellar neurons as it does in the hippocampus. In cerebellar slices, when L-type channels are blocked, mIPSCs in MLIs and PCs decrease in frequency; conversely, when L-type channels are activated, mIPSCs in MLIs and PCs increase in frequency (Rey et al., 2020). Since Ca_V_1.3 is the major L-type channel expressed in MLIs (Kozareva et al., 2021), it is likely the presynaptic L-type channel that modulates cerebellar MLI GABA release.

Taken together these data suggest that Ca_V_1.3 does not alter major aspects of cerebellar anatomy. Instead, our data suggest that Ca_V_1.3 plays an important role in PC structure, which may contribute to some of the behavioral phenotypes observed in Ca_V_1.3 KO mice (Lauffer et al., 2022), although it is notable that we do not observe deficits in eyeblink conditioning. We expected loss of PC complexity to impact eyeblink conditioning, so we were surprised to observe normal acquisition, timing, and expression of delay EBC. However, there are several examples of the converse, that is, mouse models of autism in which delay EBC is impaired but PC structure appears normal (Kloth et al., 2015). Global loss of Ca_V_1.3 in mice impairs cognitive function in several systems including development of fear memories and addictive behaviors, working memory and associative memory, and motor learning (Berger and Bartsch, 2014; Busquet et al., 2010; Jelitai et al., 2016; Kim et al., 2017; Lauffer et al., 2022; Marschallinger et al., 2015; McKinney and Murphy, 2006). Previous work with Ca_V_1.3 KO mice identified impaired consolidation of contextual fear conditioning, though cued fear conditioning was not measured (McKinney and Murphy, 2006) as Ca_V_1.3 KO mice are congenitally deaf (Eckrich et al., 2019; Hirtz et al., 2011; Jensen and Mody, 2001; Platzer et al., 2000). Prior work from the Williams lab found that Ca_V_1.3 KO mice display no changes in motor activity via the open field test but do have impairments in locomotor adaptation and learning as measured by the accelerating rotarod and Erasmus ladder (Lauffer et al., 2022). On the Erasmus ladder, Ca_V_1.3 KO mice display no motor coordination deficits as measured by missteps (Lauffer et al., 2022) but do have an impairment in gait adaptation, which is a task linked to cerebellar function (Vinueza Veloz et al., 2015). Experiments in PC-specific conditional knockout mice would be required to know how Ca_V_1.3 expression in PCs affects specific behaviors. Interestingly, loss of the non-canonical Wnt signaling protein PRICKLE2 reduces Purkinje cell excitability without affecting eyeblink conditioning (Abbott et al., 2024), suggesting that perhaps larger disturbances of PC activity and structure are required to impact this form of associative learning.

One unanticipated outcome of this work is our findings regarding the variability of cerebellar anatomy across lobules of the vermis. While each lobule contains largely similar neuronal types (n.b. unipolar brush cells which are predominantly observed in lobules IX and X) and cortical layer structure, we found that multiple parameters vary across lobules, including granule and molecular layer thickness, numbers of MLIs, and both PC spine density and spine type. Different genetic strains of mice are known to display differences in cerebellar foliation as well (Inouye and Oda, 1980; Neumann et al., 1990). Our data may serve as a reference for further work in this specific genetic background (C57BL6/NTac) and highlight the importance of using comparisons that are matched for genetic strain and lobule in structural analyses.

## Declarations

Funding: This work was funded by KL2TR002536 (AJW), the Roy J. Carver Charitable Trust (AJW and SH), NS104836 (SH), NINDS T32NS007124 (AJK), the Summer Undergraduate Research Program at the University of Iowa (JBM), the Biomedical Scholars Summer Undergraduate Research Program (MP), and the iDREAM Program (MFHA). This work utilized the Leica LMD7000 in the University of Iowa Central Microscopy Research Facilities that was purchased with funding from the NIH SIG grant 1 S10 OD016316-01. Thanks to Jordan Samuel and Hsiang Wen for technical assistance, and to John Freeman, Hunter Halverson, and members of the Williams lab for feedback on the manuscript. The funders had no role in study design, data collection and analysis, or preparation of the manuscript.

